# Genomics and CT imaging reveal diversity in silk genes and gland morphology of webspinners

**DOI:** 10.64898/2026.08.07.743568

**Authors:** Amanda Markee, Lillian J. Buck, Daniel D. Davis, Janice S. Edgerly, Edward L. Stanley, Jessica L. Ware, Akito Y. Kawahara, Ashlyn Powell, Cheryl Y. Hayashi, Richard H. Baker, Paul B. Frandsen

## Abstract

Webspinners (Insecta: Embioptera) are an unusual insect order that are known for their subsocial behavior and prolific silk-production. Due to their unique foreleg silk glands, and spider-like ability to produce silk throughout their entire life cycle, webspinners are hypothesized to have evolved silk independently from other arthropod lineages. To date, there are no reference-quality genomes available for the order, preventing the study of their silk gene origination and diversification. Here, we assembled PacBio HiFi reference genomes and characterized the silk genes present in two webspinner species, *Aposthonia ceylonica* and *Oligotoma nigra*. The genomes reveal multiple full-length copies of the primary Embioptera silk gene*, e-fibroin,* that have undergone both ancestral and recent gene duplications within the group. For both species, all *e-fibroin* paralogs show the presence of complex repeat units consisting of multiple exons and introns that are remarkably homogenized across each gene. We also used *μ*CT-scanning of the internal silk glands to provide details concerning the localization of silk production in foreleg tarsi, and interspecific morphology.

**Article summary:** This study introduces the first high-quality genomes for webspinners, enabling new research on silk for evolutionary biologists and materials scientists alike. The authors sequenced two embiopteran species, *Aposthonia ceylonica* and *Oligotoma nigra*, to compare silk genes and gland structure using micro-computed tomography, an imaging method that shows internal anatomy in detail. They found multiple copies of the primary silk gene in both species that likely arose from multiple duplication events at different evolutionary times. These silk genes exhibit unusual gene structure with hierarchically organized repeat units that are highly homogenized within a gene. The findings show that silk genes have a complex evolutionary history in webspinners and provide a foundation for studying silk diversity within the order, and in the broader context of insect silk.

## Introduction

Embioptera insects (webspinners) consist of approximately 400 described species, all of which produce silk that is essential to their survival and overall fitness (Ross 2001; Ross 2003a; Ross 2003b; Szumik et al. 2008). Webspinners typically live communally in subsocial colonies, with most species residing in the tropics feeding primarily on lichen and epiphytic algae. Other species are detritivores living in more arid environments, and feed on decaying plant matter such as leaf litter (Ross 2000). These colonies are made up of silk-constructed domiciles, known as silk galleries, which provide protection from predators and shelter against rain (Edgerly 1994; Barber et al. 2025) (Figure 1). Due to their silk’s unique properties when interacting with water, and its extraordinary fineness compared to other insect silks (Stokes et al. 2018), webspinners have been the focus of several studies in silk gland morphology (Collin et al. 2009; Büsse et al. 2015; Büsse et al. 2016), structural protein characterization (Addison et al. 2014; Barber et al. 2025), and hydrophobicity (Stokes et al. 2018; Harper et al. 2021) to characterize the variation in silk phenotype across the order.

**Figure 1.**
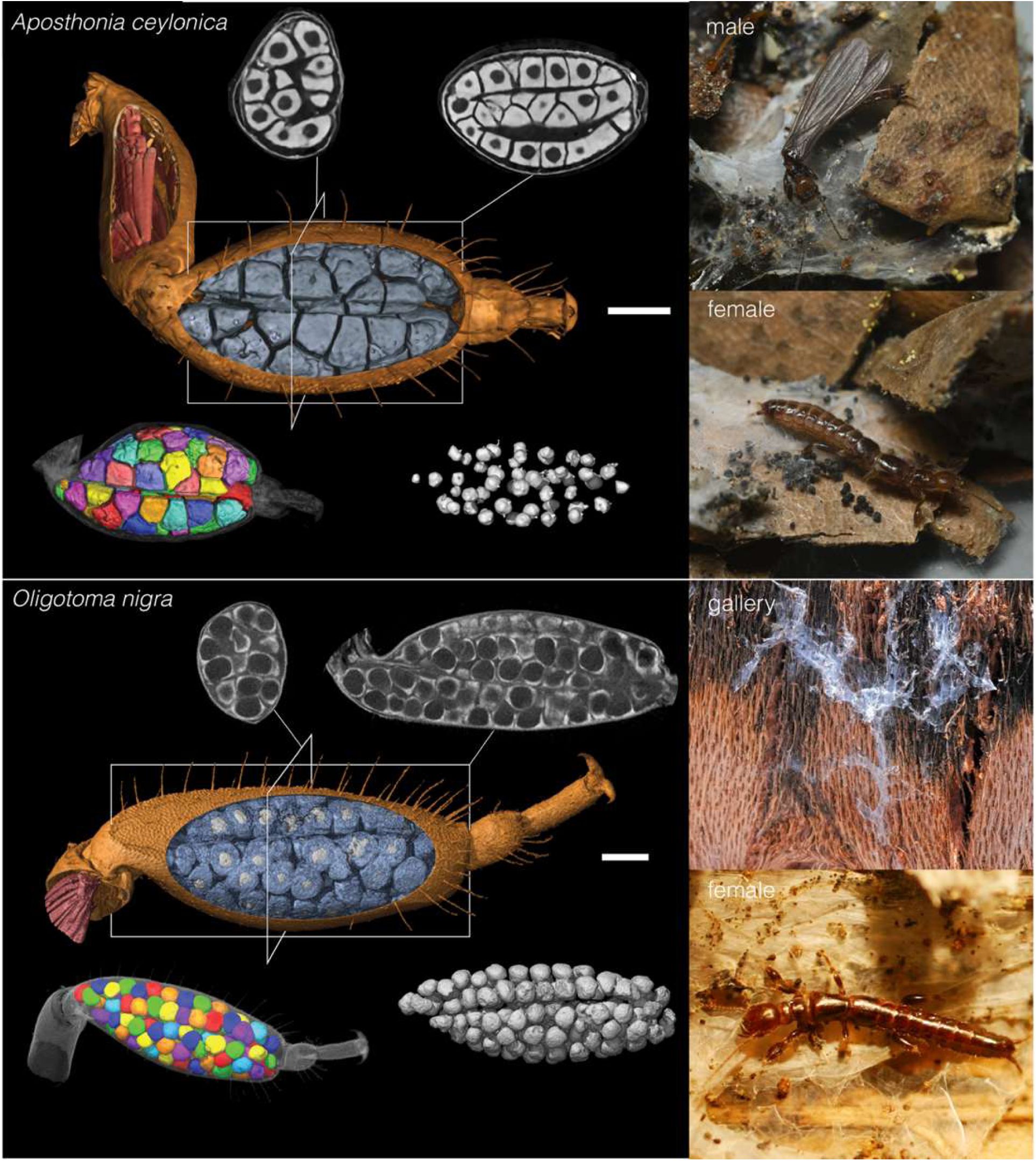
Silk spinning morphology and behavior of Embioptera. (Top) *Aposthonia ceylonica* with representative photographs of winged adult male and wingless adult female outside of their silk galleries. (Bottom) *Oligotoma nigra* representative photographs of the silk gallery and wingless adult female. MicroCT scans of silk-producing organs found within tarsi, including multiple silk glands and reservoirs produced in this study to provide morphological differences between species. Scalebars are 0.1 mm. Photos of *A.ceylonica* by C. Riley Nelson. Photos of *O. nigra* by Janice S. Edgerly.

Silk spinning morphologies exhibit substantial diversity across arthropods. For example, moths produce silk from a single pair of modified labial (salivary) glands, while spiders synthesize multiple silk types from a suite of specialized abdominal glands (Hayashi and Lewis 2001; Sehnal and Sutherland 2008). In contrast to other insects, webspinners possess silk glands in their forelimbs and, like spiders, retain the ability to produce silk throughout their entire life cycle (Edgerly et al. 2012; Büsse et al. 2015; Büsse 2019). Within each foreleg, several separate silk glands fill the inner cavity-space of the basotarsomere, with each gland composed of multinucleate secretory cells, glandular tissue, a protein reservoir, and a hollowed, hair-like silk ejector (Figure 1) (Mukerji 1927 Nov 30; Büsse et al. 2016). Liquid silk proteins are sequestered from the glandular tissue and stored in the reservoir, and then as needed, are spun into usable fibers through the duct and silk ejectors after protein rearrangement. Büsse and co-authors (2019) discovered the unique spinning mechanism in Embioptera, where adhesion discs attach to the substrate that silk ejectors press against, creating a shearing effect on the fiber. As each silk ejector is connected to an individual gland, a single embiopteran can contain hundreds of silk glands in their tarsi (Ross 2000; Collin et al. 2009).

Variation in silk gland morphology (Sehnal and Sutherland 2008), silk spinning behaviors (McMillan et al. 2016), biotic factors (colony density, and prey size) (Barve et al. 2023), and abiotic factors (temperature, humidity, and annual precipitation) (Edgerly et al. 2020; Shenoy et al. 2020) are all thought to impact Embioptera silk phenotype. Previous studies (Büsse et al. 2019; Edgerly et al., 2020) suggest that differences in spinning behaviors and silk morphologies are associated with animal body size, with larger-bodied individuals exhibiting more complex spinning behaviors and greater quantities of silk. This observed correlation, termed The Body Size Hypothesis (Busse et al. 2019; Edgerly et al., 2020), may reflect increased ecological demands for silk due to larger individuals and colonies. However, this has not yet to be explored widely and has not considered how the genetic basis for silk may play into these correlations.

Outside of model lepidopterans and spiders, the relationship between silk genotype and phenotype is still not well-established. To date, the only genetic data available for many embiopteran species is limited to single loci for phylogenetic analyses or partial silk-gene sequences, with no existing reference genome available for the order (Collin et al. 2011; Harper et al. 2021). The most recent comprehensive characterization for silk genes in this order (Harper et al. 2021) generated partial transcripts using mRNA of silk glands for two tropical species (*Antipaluria urichi* and *Pararhagadochir trinitatis).* However, due to the length and repetitive nature of silk genes, these transcripts were fragmented and only represent a single gene-copy, making genomic inference related to silk production challenging. Collin et al. 2009 provides an analysis the silk gene of *Antipaluria urichi* using cDNA finding that the fibroin was primarily composed of glycine, serine, and alanine in a repetitive GS or GA structure with repetitive groupings of codons. The absence of a reference genome, and sequence availability for broader taxonomic sampling, has prevented the complete characterization of the long and repeat-rich silk genes in this group.

Silk has evolved multiple times across arthropods (Sutherland et al. 2010). Although derived independently, the genes that encode primary silk proteins (fibroins) often have structural similarities, such as a glycine-rich repetitive core, flanked by non-repetitive N- and C-terminal domains that serve important roles in fiber formation and protein interactions (Ayoub et al. 2007; Rising and Johansson 2015). In the most well-studied insect silks, namely Lepidoptera and their sister-group, Trichoptera, the typical intron-exon structure for the silk gene *h-fibroin* is relatively simple: one short exon containing the N-terminus, followed by a single short intron and a second, long exon that contains a central repetitive region and C-terminus (Zhou et al. 2000; Zhang et al. 2024; Standring et al. 2025). There is a substantial body of research on the relationship between gene characteristics, such as repeat motif structure and amino acid composition, and mechanical performance of lepidopteran silks, due to model organisms like the domestic silkworm (*Bombyx mori*). However, outside of Holometabola, there are currently no annotated full-length insect *fibroins* characterized, leaving a significant knowledge gap in our understanding of non-model insect silks.

To investigate the diversity and evolutionary history of silk traits in Embioptera, we generated high-quality reference genome assemblies for two webspinner species (Family: Oligotomidae), *Aposthonia ceylonica* and *Oligotoma nigra*, and identified the primary silk genes, termed “embiopteran fibroin” or *e-fibroin*, within these genomes. We also use computed tomography (CT) scanning to construct 3D models for assessing differences in silk spinning morphology between both species. *Aposthonia ceylonica* is an arboreal species native to tropical Africa and Asia that experiences intense rainfall in its native environment and has been the focus of numerous studies evaluating behavior and silk qualities (Auletta 2012; Edgerly et al. 2012; Stokes et al. 2018). *Oligotoma nigra* is native to Afro-Eurasia and has been introduced to parts of North America (Büsse et al. 2015). As an opportunistic species, they can be found both arboreally in the crevices of tree bark, as well as terrestrially in detritus, depending on environmental conditions like temperature and humidity (Büsse et al. 2015; Edgerly et al. 2020). These species serve as an ideal system for evaluating interspecific and intraspecific silk variation due to their diverse ecological niches for silk use, and previously available work on silk properties in both groups (Edgerly et al. 2012; Barber et al. 2025). By providing a comprehensive genomic characterization of *e-fibroins*, we can identify structural differences that may be associated with differing habitats and silk usage between these species. In addition, based on body-size differences and habitation of different niches, we expect variation in the number of silk glands present. Analysis of genomic features in tandem to characterizing morphological diversity presents an opportunity to better understand silk composition for functional relevance in a pair of closely related taxa. *O. nigra* is also abundant, as an introduced species to the United States, allowing for further investigation into intraspecific differences in silk properties.

## Results

### μCT Scanning of Silk Glands

Both *Aposthonia ceylonica* and *Oligotoma nigra* possess numerous silk glands enclosed within the basal segment of their foreleg tarsi. Each gland is connected to an individual duct and silk ejector (Figure 1). Previous studies estimated the number of silk glands in other embiopteran species by microscopically counting external silk-ejectors, assuming a one-to-one correspondence between ejectors and glands. Building on the studies of Embioptera spinning morphology (Edgerly et al. 2012; Büsse et al. 2015), we used 3D reconstructed models to determine the number of silk glands present in each species. We find that *A. ceylonica* and *O. nigra* have 52 and 112 silk glands per tarsus, respectively, values that are consistent with earlier estimates for both species.

### Whole Genome Sequencing, Assembly and Annotation

From our HiFi DNA library for *Aposthonia ceylonica,* we sequenced 95.2 Gbp across 7.4 million reads with a read N50 of 16.1 kbp and an average read length of 13.5 kbp. Genome-size and heterozygosity estimations using k-mer count (Kokot et al. 2017) (KMC; RRID:SCR_001245) and GenomeScope 2.0 (Ranallo-Benavidez et al. 2020) predicted a genome length of ∼2.3 Gbp, with 2.59% heterozygosity and 41x coverage. For *Oligotoma nigra*, we sequenced 85.7 Gbp of HiFi sequence data across 5.2 million reads—with a read N50 of 17.4 kbp and an average read length of 16.4 kbp. Genome-size estimations and heterozygosity were predicted with the same techniques that we used for *A. ceylonica* and resulted in a genome length of ∼2.6 Gbp, 1.93% heterozygosity and 33x coverage. The findings for both species are consistent with genome size estimations predicted by flow cytometry for other species across the order (Kelly et al. 2024).

The initial *A. ceylonica* assembly contained 3.97 Gbp spread across 1028 contigs, with a contig L50 of 103, a contig N50 of 10.4 Mbp, and gene completeness of 98.68% (60.13% single-copy; 38.55% duplicated) (Supplemental Table 1). Purge_dups analysis identified 376 contigs as haplotypic duplicates, which were subsequently removed. Contaminant contigs (N=53) were identified using BlobTools v1.1.1 (Laetsch et al. 2017; Laetsch and Blaxter 2017) and NCBI’s Foreign Contamination Screen (Astashyn et al. 2024), which assigned contaminants to the following phyla: Microsporidia, Streptophyta, Chordata, Pseudomonadota, and Ascomyota. After removing haplotypic duplication and contaminants, the final assembly was 3.08 Gbp in length across 620 contigs, with an L50 of 69 contigs, an N50 of 12.61 Mbp, and a final gene completeness score of 98.76% (93.93% single-copy; 4.83% duplicated) (Table 1). The initial *O. nigra* assembly prior to decontamination contained 4.01 Gbp spread across 2,833 contigs with an L50 of 111 contigs and an N50 of 10.5 Mbp (Supplemental Table 1). The gene completeness was 99.20% (78.35% single-copy; 20.85% duplicated) (Supplemental Table 1). Purge_dups removed 1,889 haplotypic duplicates. Contaminant contigs (N=16) were identified as phyla: Pseudomonadota, Bacilota, Chordata, and Streptophyta. After removing contaminants using BlobTools and removing duplication, the final assembly was 3.17 Gbp in length across 929 contigs with a final gene completeness score of 98.61% (96.34% single-copy; 2.27% duplicated) (Table 1). The final assembly lengths were larger than those estimated by GenomeScope. We attribute this to kmer-based methods underestimating repeat content as shown in other systems (Pflug et al. 2020).

**Table 1.** Genome assembly statistics and gene completeness after running purge_dups on assemblies and removing contaminant contigs. The BUSCO database used was insecta_odb10 which contains 1367 genes.

|  |  | *Aposthonia ceylonica* | *Oligotoma nigra* |
| --- | --- | --- | --- |
| Sequencing statistics | Sequencing reads (Gbp) | 95.20 | 85.64 |
| Final assembly statistics | Contigs | 598 | 929 |
|  | L50 | 69 | 76 |
|  | N50 (Mbp) | 12.6 | 12.1 |
|  | Total length (Gbp) | 3.08 | 3.17 |
| Compleasm | Complete (%) | 98.76 | 98.61 |
|  | Single-copy (%) | 93.93 | 96.34 |
|  | Duplicated (%) | 4.83 | 2.27 |
|  | Fragmented (%) | 0.22 | 0.22 |
|  | Missing (%) | 1.02 | 1.17 |
| Sequencing statistics | Sequencing reads (Gbp) | 95.20 | 85.64 |
| Annotation | Program | Helixer | Augustus |
|  | Putative genes | 44,691 | 52,790 |
|  | Complete (%) | 84.13 | 97.88 |
|  | Single-copy (%) | 67.52 | 73.45 |
|  | Duplicated (%) | 16.61 | 24.43 |

Our genome size estimation from GenomeScope predicted that 56.2% of the *A. ceylonica* genome was comprised of repetitive elements, while 45.6% of the *O. nigra* was estimated to be repetitive. After Earl Grey (Baril et al. 2024) repeat modeling and masking, we classified 77.6% of the *A. ceylonica* genome assembly as repetitive and 74.53% of the *O. nigra* genome assembly as repetitive. This disparity between approaches explains the GenomeScope underestimates of genome size (Supplemental Fig. S1). In both genome assemblies, most repetitive elements were unclassified, which is perhaps unsurprising, given the lack of genomic resources for this group and their lack of representation in the repeat libraries (Sproul et al. 2023) (Supplemental Fig. S2-S3). We used multiple genome annotation analyses for both genomes and chose the best annotation for each based on compleasm protein scores. For the *A. ceylonica* genome, 44,691 coding sequences were identified with Helixer v0.3.4 (Stiehler et al. 2021; Holst et al. 2023) via the Helixer web server (https://www.plabipd.de/helixer_main.html). Of these genes, 80% had BLAST hits against the NCBI non-redundant protein database, 61% received GO Mappings, and 32% were functionally annotated with Blast2GO. The protein compleasm results indicated that the final annotation was 84.13% complete (67.52% single-copy; 16.61% duplicated). For the *O. nigra* genome, the Augustus whole-genome annotation yielded 52,790 amino acid coding sequences with the following annotation quality statistics as calculated by compleasm v0.2.5: 97.88% complete (73.45% single-copy; 24.43% duplicated). Of the genes, 65% had BLAST hits against the NCBI protein database, 43% received GO Mappings, and 20% were functionally annotated with Blast2GO.

### Embioptera fibroins (e-fibroin) have undergone extensive gene duplication

In contrast to other large repetitive insect silk genes such as the heavy chain fibroin (*h-fibroin)* of Lepidoptera and Trichoptera, which is typically single-copy, the genome assemblies of both embiopteran species contained multiple copies of *e-fibroin*. We identified putative signal peptides in each paralog and further verified the annotation of these genes as encoding silk proteins by mapping RNA reads derived from silk glands to each putative *e-fibroin.* In Iso-Seq sequencing of pooled silk gland samples from *A. ceylonica*, 36-51% of all transcripts mapped to the putative *e-fibroins*, compared to 0.01-0.02% of Iso-Seq transcripts in whole body specimens. This difference in transcript count strongly supports that these are silk genes. Four paralogous copies of *e-fibroin* were recovered in the *A. ceylonica* genome, and five copies were recovered in *O. nigra. A. ceylonica e-fibroin* genes were tandemly arrayed in the same orientation in a cluster spanning 543.5 kbp and with no other intervening genes (Figure 2a). Four of the *O. nigra e-fibroin* genes were similarly located in a cluster, with one inverted copy of the gene separated by approximately 1.1 Mbp on the same contig. To ensure paralogs were accurately represented, and not the result of assembly errors that incorrectly inserted reads from a single paralog into multiple locations, we compared coverage distribution between paralogs and background genes. We find equal coverage distribution across all paralogs compared to other genes present on their respective contigs and the genome as a whole (Supplemental Fig. S4).

**Figure 2a.**
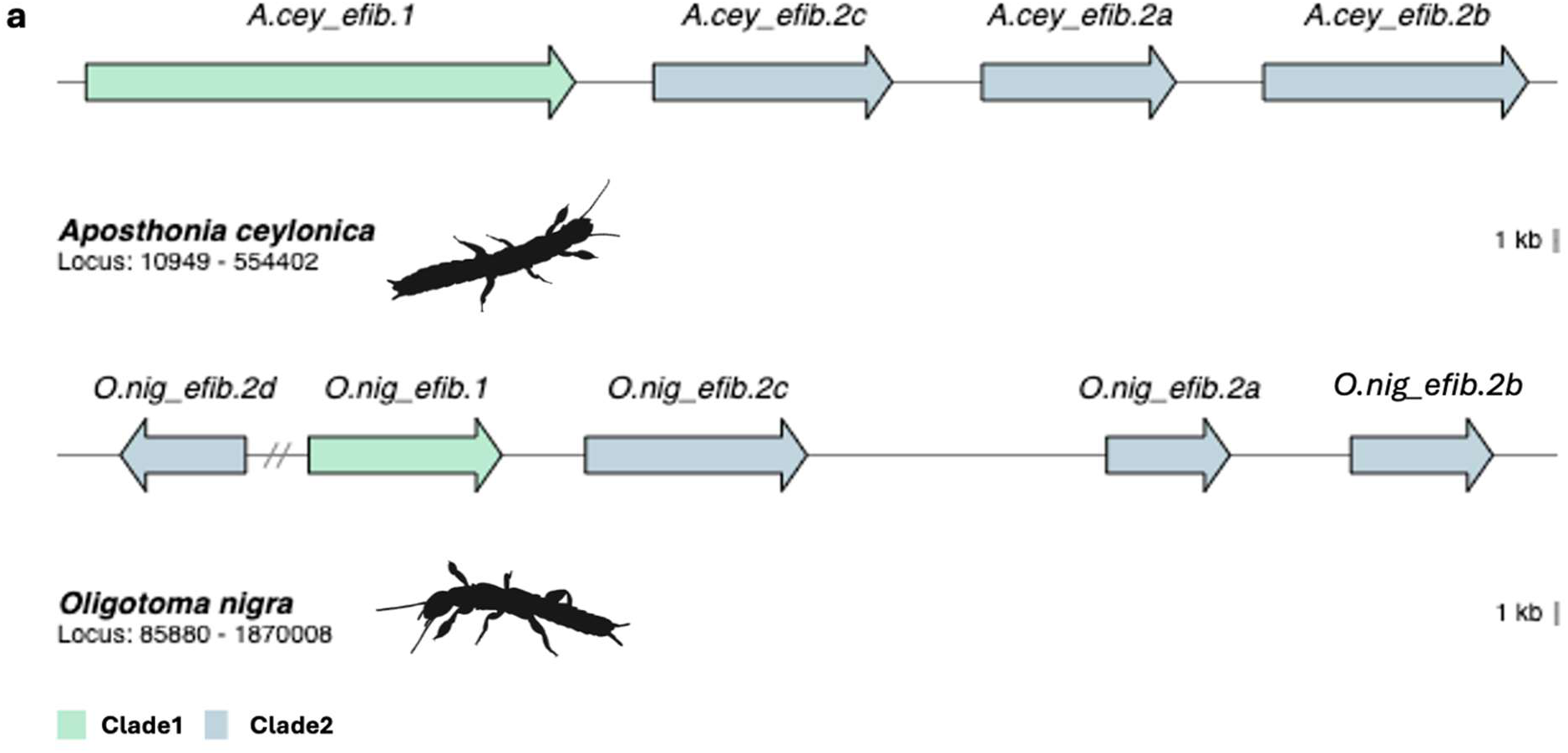
Embioptera *e-fibroin* genomic location and evolution. a) Organization of *e-fibroin* gene clusters on their respective contigs. Hash marks in *Oligotoma nigra* represent separation across the contig. Gene colors correspond to their clade placement in the gene family tree (Figure 2b).

Phylogenetic analysis of the inferred amino-(N) and carboxy (C)-terminal protein sequences of *e-fibroins* (Figure 2b), combined with previously published partial sequences from a few other species (GenBank accessions MW698943.1, MW698944.1, EU170437.1, HM189214.1, HM189215.1) reveals two primary clades containing *e-fibroin* genes from both *A. ceylonica* and *O. nigra,* suggesting that there was a duplication of the *e-fibroin* gene prior to the split between the two species (Figure 2b). Following this initial duplication, it is likely that subsequent duplications occurred within each species (Figure 2b). Four of the *O. nigra* genes were recovered as monophyletic on the tree and therefore, are consistent with recent duplications. The three *A. ceylonica* genes that are sister to this *O. nigra* group are paraphyletic, but this may reflect weak phylogenetic support due to the short nature of the terminal domains rather than a series of losses in *O. nigra.* Clade 1 includes two additional species (*S. davisi* and *Archembia sp.)*, suggesting they may also have multiple *e-fibroin* paralogs (Figure 2b). While the bootstrap values separating Clade1 from Clade2 are low, this is driven primarily by the Genbank taxa that lack N-termini sequences. A phylogenetic analysis of just the *A. ceylonica* and *O. nigra* paralogs separates the Clade1 taxa from Clade2 with a bootstrap of 98.6 (Supplemental Fig. S5).

**Figure 2b-2c.**
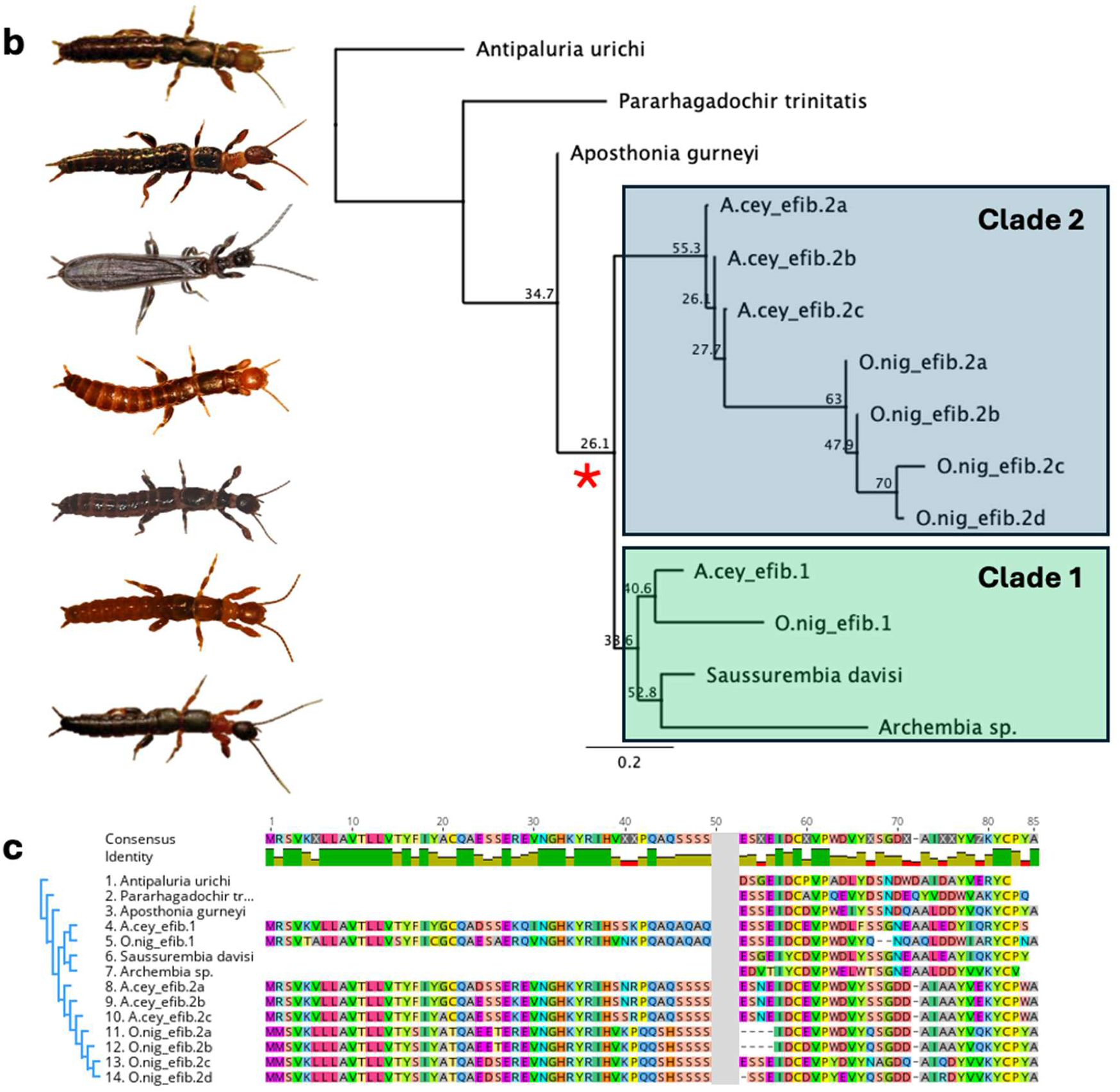
Embioptera *e-fibroin* genomic location and evolution. b) Phylogeny of previously published *e-fibroin* termini, and those generated from this study. Red asterisk indicates the ancestral duplication of the *e-fibroin* gene. The tree is rooted with the *Antipaluria urichi e-fibroin* as this is the most distantly related embiopteran species with an available terminal sequence. Habitus photos of each species in the tree are shown to the left in the order in which their sequences appear, starting at the root: *Antipaluria urichi, Pararhagadochir trinitatis, Aposthonia gurneyi, Aposthonia ceylonica, Oligotoma nigra, Saussurembia davisi, Archembia sp*. Photos courtesy of Janice Edgerly, Edward Rooks, and iNaturalist contributor Ella Sanctuary. c) Concatenated N- and C-terminal region alignment used in tree reconstruction. The gray block separates the termini.

### Atypical insect fibroin organization: multiexonic with complex repeats

Unlike the *h-fibroin* genes in most Lepidoptera, which have only one intron, all *e-fibroin* sequences recovered from both species have a highly complex intron-exon structure involving many small repetitive exons (Figure 3, Supplemental Fig. S6). On average, *A. ceylonica e-fibroin* paralogs contain 136 exons (range: 96-196), while *O. nigra e-fibroin* paralogs contain 72 exons on average (range: 33-133). In general, the introns within each gene can be classified into multiple groups based on their sequence identity and are organized in regularly spaced patterns throughout the gene. (Supplemental Fig. S7). For instance, *A.cey_efib.2c* contains three primary intron types that are 96.57% identical among all copies within a type, but less than 50% identity between types (Figure 3). Members of each intron type occur in a regular pattern (Type1-Type2-Type3) that repeat throughout the gene, suggesting that a sequence unit comprised of three exons and three introns represents the primary repeat unit. These large repeat units (which we refer to as Ensemble Repeats, ERs) span approximately 2.2 kbp across the three exons and introns in *A.cey_efib.2c* and have an average percent identity at the nucleotide level of 95.9%, when excluding large indels. In addition, the introns from different paralogs within each species are divergent from one another, with an average identity of 27% when the consensus sequences of each intron type for a given gene ER are compared to the consensus sequences of other intron types from different *e-fibroin* paralogs in the same species (Supplemental Table 2). Among the nine *e-fibroin* sequences identified in these two genomes, all but one include multi-exonic ER units with two genes having an ER consisting of 2 introns and exons, three genes having an ER of 3 introns and exons, two genes with an ER with 4 units, and one with 5 (Figure 3). This highly homogenized intron-exon structure of ER units with multiple exons has previously been observed in spider silk genes (Baker et al. 2025) but not yet in insect silk genes.

**Figure 3.**
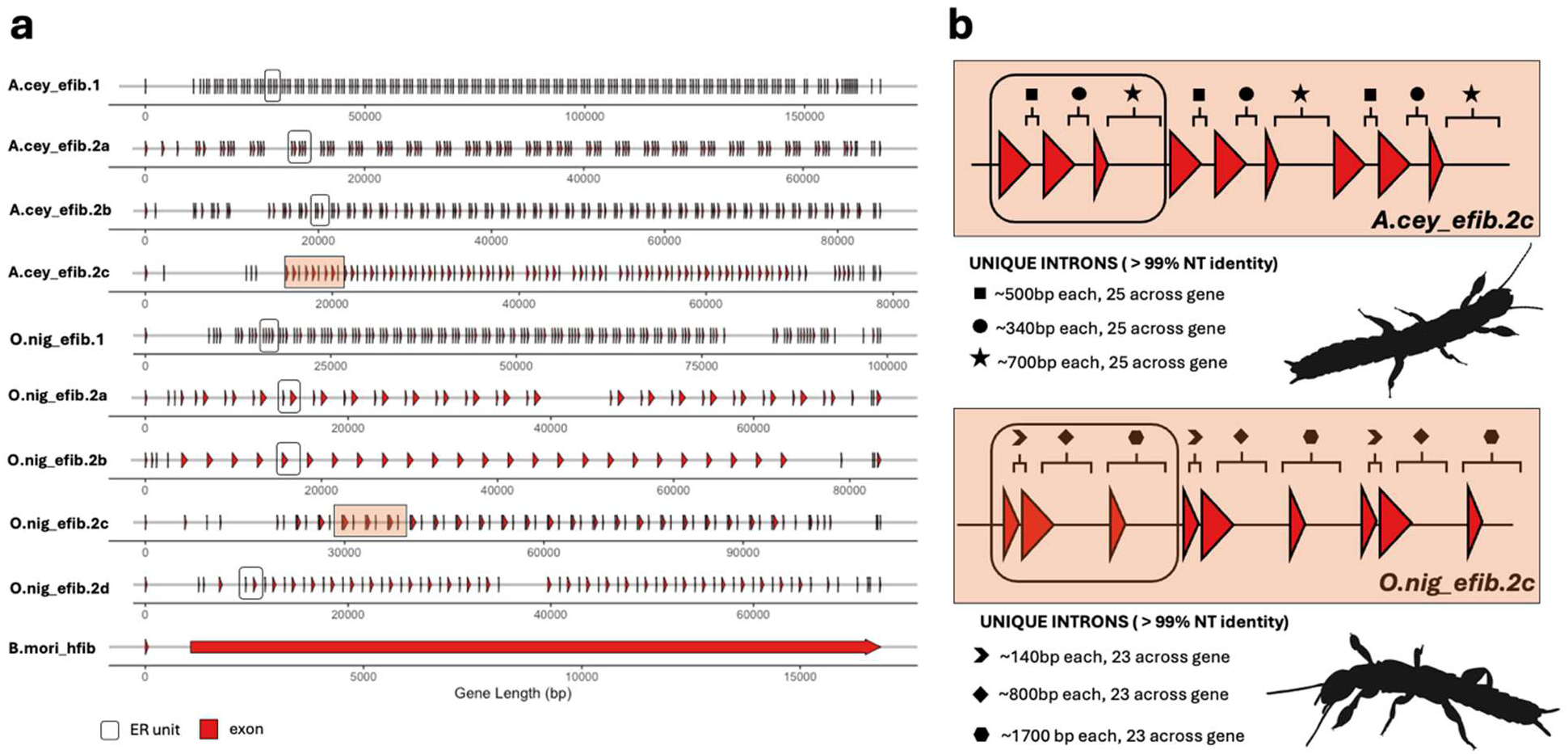
Intron-exon structure of insect *fibroin* genes. a) *Aposthonia ceylonica* and *Oligotoma nigra e-fibroin* silk genes have many introns and exons compared to the *h-fibroin* of the domestic silkworm, *Bombyx mori*, which includes only one intron and two exons. Red triangles along the X-axis (contig) indicate exons, while the black rectangular outlines surrounding a group of red triangles represent examples of ensemble repeat (ER) units b) Examples of the ER structure of *e-fibroin*. Shaded boxes in *A.cey_efib.2c* and *O.nig_efib.2a* highlight several ER units depicted in more detail, highlighting both the multiexonic patterns, and the homogenization of introns.

### Repeats units are highly homogenized for simple amino acid motifs

Despite the extensive gene duplication of *e-fibroins* in *A. ceylonica* and *O. nigra*, the repetitive protein sequences of the paralogs are similar to each other in amino acid sequence and composition. All nine genes are dominated by simple alternating glycine and serine (GS), or glycine and alanine (GA) motifs, with the three amino acids comprising, on average, 94.4% of the protein sequence in all genes (range: 93% - 96.8%). The GA motifs are common (43.1%) in Clade1 genes but are not prevalent (6.7%) in Clade2 genes. Within Clade2, GA motifs appear in all *O.nigra* ERs, but only in half of *A.ceylonca* ERs (Supplemental Fig. S8). Each gene also contains atypical amino acids that generally occur as a single residue interspersed among the GS/GA repeat motifs (Figure 4a). These uncharacteristic residues correspond to the pattern found in a previous study of partial *Antipaluria urichi e-fibroin* sequence (Harper et al. 2021). The most common of these residues include asparagine, threonine, and glutamate (polar residues), as well as glutamic acid (acidic and charged) (Figure 4c), but there are notable differences in the occurrence of these residues among the *e-fibroin* paralogs.

**Figure 4.**
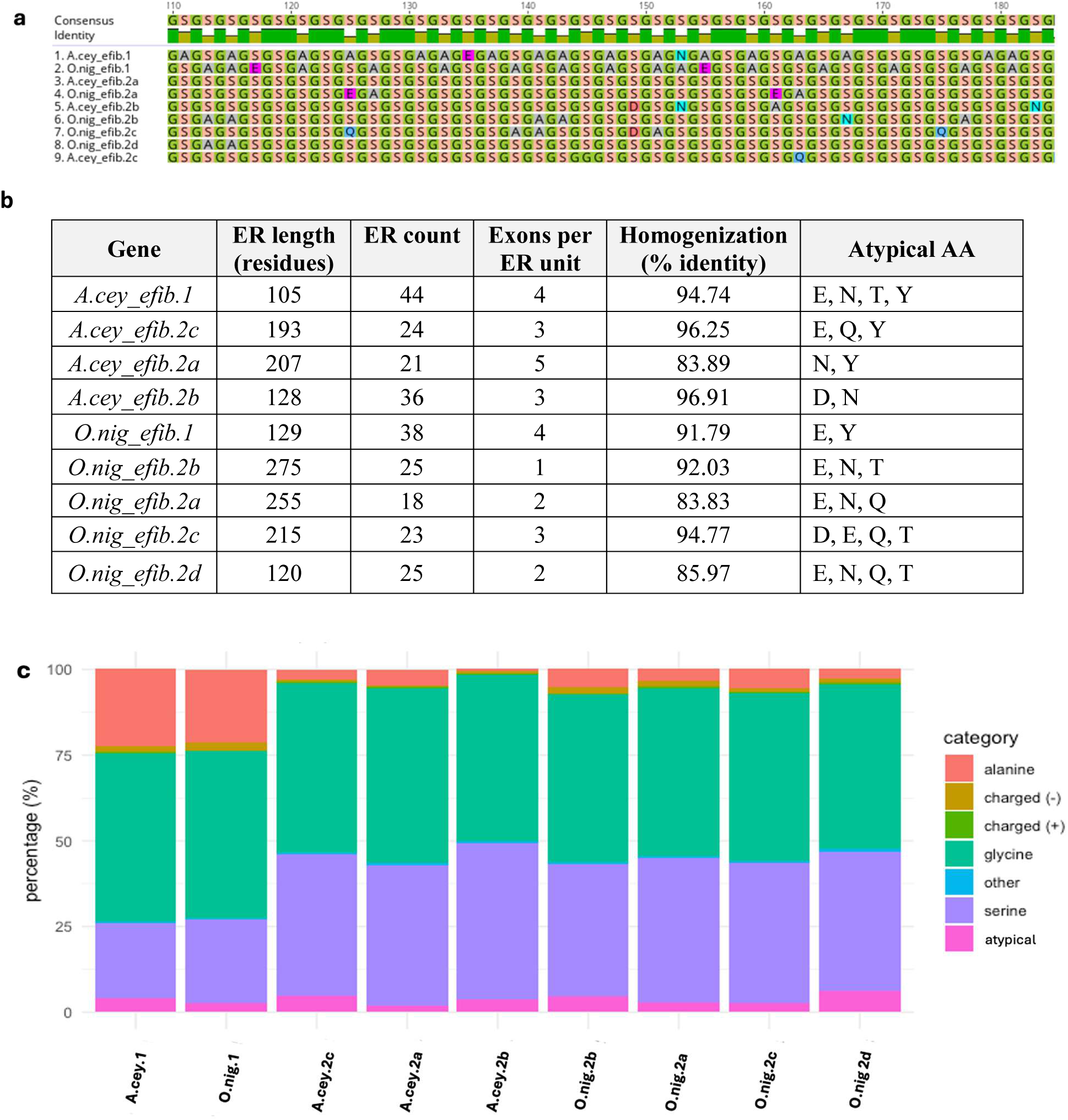
Characteristics of the nine e-fibroin predicted proteins. a) A 62 amino-acid sequence region from the protein alignment of nine *e-fibroin* sequences ensemble repeat units, showing high sequence similarity with glycine and serine richness, but differentially spaced atypical amino acids in between. b) Ensemble repeat (ER) features and length statistics, showing highly homogenized sequence identities, ER abundance and presence of atypical amino acids. c) E-fibroin amino acid compositions. “Charged (−)” consists of amino acid residues D and E; “charged (+)” consists of residues H, R and K; “atypical” consists of Q, N, T and Y; and “other” consists of all remaining residues.

One unifying characteristic across all *e-fibroin* genes in *A. ceylonica* and *O. nigra* is the homogenization of ER units. At the peptide level, a majority of ensemble repeats consistently show over 90% shared sequence identity, suggesting that the genetic mechanism that conserves repeat units likely prioritizes repeat identity through the units evolving together (Figure 4b). We found three primary factors that influence variation in the homogenization and percent identity of ER units: the insertion of indels with exons, nucleotide variation, and/or the addition of extra exons within the ER. Variation in these three instances will disrupt the identical nature of ERs, and lead to lower homogenization indices. Full consensus ensemble repeats for each gene can be found in Supplemental Fig. S8.

### Allelic variation: length disparity of protein-coding regions within individuals

For each *e-fibroin* of *A. ceylonica*, we recovered extensive allelic variation, highlighting another layer of genetic diversity (Table 2). Following similar observed patterns of silk-gene allelic variation in spiders, caddisflies, and moths (Frandsen et al. 2023), we attribute a large majority of this variation to the presence of large indels across the gene (Supplemental Fig. S9), resulting in substantial allelic differences in the number of exons, hence total length, within a single gene. In *A. ceylonica,* protein coding length variation ranges between 111bp – 2313bp across all *e-fibroin* paralogs. Due to fragmentation in the alternate assembly of *O. nigra,* allelic diversity and length variation in the alternate haplotype was not assessed.

**Table 2.** Gene and coding sequence (CDS) lengths and exon numbers for *e-fibroin* paralogs in *A. ceylonica* and *O. nigra*. The upper section of the table depicts results for *A. ceylonica.* The lower section depicts the results for *O. nigra*.

| *Aposthonia ceylonica* |  |  |  |  |
| --- | --- | --- | --- | --- |
| Gene | Haplotype | Gene length (bp) | CDS length (bp) | # of exons |
| *A.cey_efib.1* | Primary | 167,339 | 15,600 | 196 |
|  | Alternate | 173,690 | 15,792 | 196 |
| *A.cey_efib.2c* | Primary | 78,682 | 15,627 | 96 |
|  | Alternate | 90,018 | 16,797 | 106 |
| *A.cey_efib.2a* | Primary | 67,088 | 14,412 | 122 |
|  | Alternate | 73,019 | 14,301 | 162 |
| *A.cey_efib.2b* | Primary | 84,975 | 15,549 | 123 |
|  | Alternate | 99,660 | 13,236 | 109 |

| *Oligotoma nigra* |  |  |  |  |
| --- | --- | --- | --- | --- |
| Gene | Haplotype | Gene length (bp) | CDS length (bp) | # of exons |
| *O.nig_efib.1* | Primary | 99,120 | 16,848 | 136 |
| *O.nig_efib.2c* | Primary | 110,726 | 17,031 | 82 |
| *O.nig_efib.2a* | Primary | 72,538 | 14,022 | 48 |
| *O.nig_efib.2b* | Primary | 83,526 | 15,459 | 33 |
| *O.nig_efib.2d* | Primary | 72,560 | 10,506 | 65 |

## Discussion

### Comparative morphology of silk-glands with μCT scanning

It has been hypothesized that larger embiopterans have a greater number of silk glands or a larger protein reservoir because they can produce more silk (Büsse et al. 2015). For example, Ross 2000 estimated that *O. nigra* has approximately 150 silk glands per tarsus, and Edgerly et al. 2012 estimated *A. ceylonica* to have approximately 53 glands. One of the largest species, *Antipaluria urichi*, has been estimated to have approximately 230 glands. *Antipaluria urichi* is 1.6x longer than *O. nigra* (Ross 1957; Edgerly et al. 2002; Collin et al. 2009) and has roughly 1.5x more silk glands. In this context, gland number appears to scale positively with body size, a patterned termed the Body Size Hypothesis (Büsse et al. 2015). While our results do not fully support this hypothesis, we recognize the limitation of our sampling, as we use two closely related species and single representatives for these analyses. Despite being relatively similar in body size, *O. nigra* has approximately twice as many silk glands as *A. ceylonica*. Thus, the relationship between body size and silk production capacity may be less straightforward than previously thought. A general trend likely exists in which larger-bodied embiopterans possess more silk glands and produce greater quantities of silk; however, ecological or life-history factors may permit specialized exceptions to this pattern. Additional *μ*CT-scanning and quantitative morphometric analyses across a broader taxonomic sampling of embiopterans are necessary to further clarify the relationship between silk production and body size.

### Numerous convergent features of Embioptera silk genes

We assembled full-length *e-fibroin* sequences in two Embioptera species revealing more detail in the molecular evolution of insect silk genes. These sequences reveal convergent characteristics to other published insect silk genes, in that they are long and highly repetitive, containing glycine-rich protein-coding regions and show conserved N- and C-terminal regions. However, in contrast to other insect silk genes, the *e-fibroins* in these species have undergone multiple gene duplications, with several paralogs discovered in each genome. The consistency in read coverage (Supplemental Fig. S4), variation in gene organization and the divergence in intron sequences across paralogs in both species (Supplemental Fig. S7) all suggest that the abundant silk gene diversity within Embioptera species did not results from assembly errors. In most Lepidoptera and Trichoptera, which have acquired silk independent from Embioptera, the *h-fibroin* gene exhibits a simple gene structure consisting of a short exon, followed by a short intron, and a large secondary exon that comprises the entire repetitive core. All the *e-fibroin* sequences, on the other hand, have complex gene organizations (Figure 3) with large repeat units comprised of several introns and exons.

Phylogenetic analyses from conserved terminal domains suggest that an ancestral duplication of the *e-fibroin* gene likely occurred prior to the divergence of *A. ceylonica* and *O. nigra* (Figure 2b). Not only is this separation of Clade1 and Clade2 supported by a high bootstrap value (Supplemental Fig S5), but the protein composition of the Clade1 genes is markedly different from the other paralogs given the increased proportion of alanine in these genes (Figure 3a). Additional lineage-specific duplications are also consistent with the recovered topology, although the precise timing and relationships among these events should be interpreted cautiously. Four *O. nigra e-fibroin* genes form a monophyletic group, consistent with relatively recent species-specific duplications. In contrast, the three *A. ceylonica* genes sister to this clade are recovered as paraphyletic. While this pattern could reflect a more complex evolutionary history, including differential gene loss or incomplete lineage sorting, it may also result from the limited phylogenetic signal available in the short terminal domains. In addition, it is possible many of the Embioptera paralogs arose before the *A. ceylonica-O. nigra* split but group together on the tree as a result of intergenic gene conversion acting on the termini sequences, as has been suggested in spiders (Sanggaard et al. 2014). Future sampling across additional embiopteran species, and analyses for detecting gene conversion are needed to distinguish between species-specific duplications, and older duplications that have undergone concerted evolution.

With respect to the abundant gene duplication and homogenized ensemble repeats, *e-fibroins* are more like spider silk fibroin genes (*spidroins*) than other insect *fibroins*. Orb-weaving spiders possess a highly diverse family of *spidroins*, with 30-40 individual genes in a single spider that can be expressed in different combinations to comprise up to eight distinct silk-types (Ayoub and Hayashi 2008; Foelix 2011). The diversification and structural organization of spidroins are closely associated with the mechanical performance of each silk-type, with different combinations of glycine, alanine and serine motifs expressed in different glandular origins (Guerette et al. 1996). More specifically, in orb-weaver dragline silks, the protein is composed of GGX, GPGXX motifs, and poly-alanine and poly-glycine-alanine repeats, with atypical amino acid residues such as asparagine, cysteine, proline, tyrosine, leucine or glutamine (Yarger et al. 2018). While molecular mechanisms and evolutionary pressures that maintain these unique gene structures are presumably variable, the presence of ensemble repeats is common among *spidroin* paralogs (Baker et al. 2022; Frandsen et al. 2023; Baker et al. 2025). These *spidroins* often contain numerous repetitive exons and introns that define the ER boundaries and introns may have evolved within these genes to facilitate the homogenization of the protein sequences within exons (Baker et al. 2022). In a few *spidroins,* ERs are comprised of two intron-exon pairs (Baker et al. 2022) but there are no known examples of spider ERs including 3 or more exons, as we observe in the *e-fibroins*. While little is known about the structure-function relationships of *e-fibroins,* their homogenized ER structure may be functionally significant by ensuring precise spacing of atypical residues. In addition, it should be noted that the species sampled in this study are from the same family and additional species within Embioptera need to be examined to determine if the extensive duplication patterns and complex gene organizations is characteristic of the group as a whole.

Both silkworm and spider silks have fairly well-understood gene structure, containing highly crystalline β-sheet regions, largely driven by extensive hydrogen bonding during β-sheet stacking. Our genomic analyses reveal that embiopteran silk, similar to many silk genes in other arthropod groups (Zhou et al. 2001; Vienneau-Hathaway et al. 2017; Zhu et al. 2026; Correa-Garhwal and Garb), are also characterized by a strong dominance of glycine–alanine (GA) and glycine–serine (GS) dipeptide motifs interspersed with highly variable spacer residues (Figure 4a). GA motifs are comparatively more hydrophobic and are predicted to favor tight β-sheet packing and crystallinity, whereas GS motifs introduce increased polarity and flexibility, potentially modulating elasticity and intermolecular interactions. We speculate that the co-occurrence of GA-rich and GS-rich motifs across different paralogous *e-fibroins* may enable fiber interactions analogous to the MaSp1/MaSp2 system in spider dragline silk. Such interactions may strike a balance of toughness and extensibility well suited for maintaining durable, yet flexible silk galleries used by subsocial colonies. While prior work has demonstrated high crystallinity in Embioptera silk (Addison et al. 2014) and highlighted mechanical similarities to silkworm silk despite its extreme fineness (Okada et al. 2008; Stokes et al. 2018), the structure–function consequences of GA/GS motif dominance, paralog diversity, and spacer variability remain unexplored in *e-fibroins.* Our genomic framework provides a foundation for testing these hypotheses and advances new insight into the molecular evolution of silk in a non-model insect lineage.

We recovered extensive protein-coding sequence length variation in *A. ceylonica e-fibroins,* consistent with a preliminary investigation of arthropod silk-gene intraspecific variation (Frandsen et al. 2023; Stewart et al. 2026). Single gene sequences of individual spiders, butterflies and caddisflies in this study show striking allelic differences within a single individual in the form of large indels interspersed throughout the protein-coding repetitive region. In spiders, it has been demonstrated that variation in repeat length contributes to the functional and mechanical properties of capture spiral silks (Li et al. 2017), but the relationship between structure and function in *fibroins* remains relatively unexplored across insect silks, especially outside of Holometabola (Stewart et al. 2026). The precise mechanisms which generate and maintain this convergent pattern across silk genes is not well established, but it is likely caused primarily by molecular processes such as unequal crossing over and replication slippage.

### Structural aspects of Embioptera silks

The strength, extensibility, and toughness of arthropod silks are determined to some extent by the amino acid motifs and composition of *fibroin* sequences (Brooks et al. 2008; Gaines IV and Marcotte Jr 2008; Savage and Gosline 2008; Creager et al. 2010; Marhabaie et al. 2014). For example, the dipeptide pattern [(GX)_N_] has been observed in the silkmoth *Bombyx mori* h-fibroins, orb-weaver spider minor ampullate spidroins, and now in Embioptera e-fibroins. Sequential GA or GS repeat motifs are thought to promote silk crystallization by facilitating the formation of ordered β-sheet domains, consistent with the central role of alanine in enhancing silk rigidity and inter-strand hydrogen bonding (Addison et al. 2014; Carrascoza Mayen et al. 2015). In contrast, atypical amino acid residues enriched in amorphous regions of silk such as tyrosine, valine, aspartic acid, and glutamic acid, disrupt GS/GA repeats, and disrupt uniform β-sheet stacking and crystalline domain formation (Vilaplana et al. 2015; Huang et al. 2023). This in turn creates flexible, elastic domains that contrast with the alanine-rich units contributing to silk’s strength.

Previous studies have hypothesized that the presence of atypical, polar amino acids may act as “spacers” to contribute to the fiber-to-film transformation that occurs in some embiopteran silks (Harper et al. 2021), where the silk changes from a string-like fiber to an amorphous film upon wetting. This occurs primarily because of the hydrophilic nature of the protein core, and the hydrophobic lipid layer surrounding the core (Addison et al. 2014; Stokes et al. 2018; Andrews et al. 2022; Barber et al. 2025). However, the functional implications of these atypical amino acids on the silk protein structure remain untested. Our results represent a mix of *e-fibroin* paralogs with a range of atypical amino acid spacers, and variation in the presence of GA/GS repeat motifs, that are not specific to either species. Generally, alanine residues occur more frequently and abundantly in *O. nigra* paralogs, with all five paralogs exhibiting GA motifs in their primary ER units. In *A. ceylonica* however, alanine is only present in half of the paralogs and occur much less, suggesting that *O. nigra* silk may exhibit higher strength and rigidity comparatively.

Consistent with our findings, Collin et al. (2011) identified conserved features across partial *e-fibroin* repeat regions from multiple taxa, including members of the genera *Saussurembia, Archembia, Haploembia, Oligotoma, Antipaluria,* and *Aposthonia*. However, their study was limited to silk-specific cDNA from the 3’ end of the transcript and only introduces a single sequence per silk gene. Interestingly, they identified a poly-serine repeat block in the repetitive region of a species sister to *Aposthonia ceylonica*, a feature absent from all ensemble repeats recovered in our *A. ceylonica* dataset. This observation highlights additional lineage-specific variation within an otherwise conserved repeat architecture. Our results extend those of Collin et al. by demonstrating that the diversity of atypical amino acid composition observed across species is also present within species, and even within a single individual possessing multiple *e-fibroin* gene copies. This previously unrecognized level of diversity in *e-fibroins* suggests that broader taxonomic sampling will be necessary to reconstruct the evolutionary history of these gene duplications. This newly discovered source of variation may have important functional implications for silk properties, and future studies will be needed to determine whether these patterns have been maintained primarily by genetic drift, natural selection, or a combination of both.

### Conclusions

Our findings show the first comprehensive annotation of full-length silk genes for the order Embioptera and reveal several striking similarities and differences with other arthropod silks. Despite occupying different ecological niches, the characteristics of the *e-fibroins* recovered in this study were similar between both species, making it difficult to discern putative functional differences from sequence data alone. Both have multiple paralogs, high degrees of complex repeat structure and homogenization, one gene represented in Clade1, multiple genes represented in Clade2, and very similar protein sequence composition. Our phylogenetic inferences from gene tree reconstruction suggest an ancestral duplication within the order giving rise to two primary *e-fibroin* clades, although the timing of this event relative to embiopteran diversification remains unresolved. More widespread sampling within Embioptera is necessary to resolve the evolutionary history of *e-fibroin* diversification, gene organization and repeat composition. In addition, future studies need to focus on the pattern of silk gene expression in order to understand the relative importance of different *e-fibroin* genes and how they are integrated into silk fibers at different developmental stages and ecological contexts.

## Materials and Methods

### Micro Computed Tomography (μCT) Scanning

One female *A. ceylonica* from a lab-reared colony was stained in 3% aqueous phosphotungstic acid for 34 days to ensure silk glands were saturated enough to discern density differences when characterizing morphology. The stained A. ceylonica was scanned at the University of Florida’s Nanoscale Research Facility (RRID:SCR_025135) on a ZEISS Versa 620 X-ray microscope (XRM) at a resolution of 904.6nm, with the following settings: X-ray voltage: 60kV, filament current: 108µA, LE1 filter, 4X objective lens, detector binning: 2, capture time: 5 seconds. One male O. nigra from a lab-reared colony was stained in a 3% phosphotungstic acid (PTA) solution for four weeks. The stained individual was mounted within melamine foam submerged in 95% ethanol and scanned at the American Museum of Natural History’s Microscopy and Imaging Facility (RRID:SCR_024760) on a ZEISS Versa 630 X-ray microscope (XRM) at a resolution of 1.078µm, with the following settings: X-ray voltage: 69.95kV, filament current: 121.47µA, LE4 filter, 4X objective lens, detector binning: 2, capture time: 10 seconds. Tomograms for both datasets were recovered using the XMReconstructor (Carl Zeiss) software suite, volumes were post-processed using Dragonfly v.2022.2 and internal morphology was segmented and reconstructed using VGStudio v.2025.1 (Volume Graphics, Heidelberg, Germany)

### DNA Extraction and Sequencing

*A. ceylonica* and *O. nigra* were reared independently in two lab captive colonies to adulthood in enclosures containing soil and leaf litter. A single male *A. ceylonica* and a single female *O. nigra* were preserved using liquid nitrogen to flash-freeze and were stored in a −80°C freezer until time of extraction. Following the manufacturer and Brigham Young University Sequencing Center’s internal protocols, high molecular weight genomic DNA was extracted from both species using the Qiagen genomic tip DNA extraction kit, then sheared to 18 Kbp fragments with a Diagenode Megaruptor (Diagenode Inc., NJ, USA). The BluePippin system (Sage Science, Beverly, MA, USA) was used to collect fractions containing >15 Kbp fragments for library preparation. Genomic libraries were prepared following the SMRTbell Express Template Prep. Kit 2.0 protocol (PacBio, Menlo Park, CA, USA). Each library was sequenced on a single flow cell for 24 movie hours with PacBio HiFi sequencing on a PacBio Revio sequencer.

### RNA (IsoSeq and Illumina Short Reads)

RNA extraction was performed for four samples of *A. ceylonica* using the Kinnex full-length RNA kit from PacBio. One sample contained tissue from one whole female, one sample contained tissue from one whole male, and two samples each contained the tarsi of 4 female individuals for a total of 8 tarsi in each sample. All libraries were run on a single SMRT cell. Size selection was not performed in order to include long transcripts. We followed the pipeline for demultiplexing, refining, and merging the SMRT cell following steps 1-3b in the Clustering CLI workflow (https://isoseq.how/clustering/cli-workflow.html). We identified putative fibroin transcripts by blasting for known *e-fibroin* termini. We then mapped these transcripts to the *A. ceylonica* genome using hisat2 v2.2.1 (Kim et al. 2019) to confirm exon boundaries. We used minimap2 v2.28 with slicing to map the transcripts to the genome to confirm intron-exon boundaries (Supplemental Fig. S6). RNA extractions for *O. nigra* was performed on three samples using a modified phenol-trizol extraction protocol. Whole bodies were prepped using the PureLink RNA Kit (Invitrogen) with bead mill homogenization and Trizol nucleic acid isolation. RNAseq reads were used primarily for identifying intron-exon boundaries across *e-fibroin* genes during manual annotation.

### Genome Assembly and Quality Control

For each species, we assembled the PacBio HiFi reads using hifiasm v.0.19.5-r587 (Cheng et al. 2021). We removed haplotypic duplication with purge_dups v.1.2.5 (Guan et al. 2020). We used Blobtools v.1.1.1 (Laetsch et al. 2017; Laetsch and Blaxter 2017) to identify contaminants based on GC content, sequencing coverage, and BLAST-based taxon identification. We removed contigs with BLAST matches to phyla other than Arthropoda from the assembly. We removed additional contigs from the *A. ceylonica* assembly using NCBI’s Foreign Contamination Screen (Astashyn et al. 2024). We measured both assembly statistics and gene completeness for 1) the initial genome assembly, 2) after the purge_dups assembly, and 3) after removing contaminants identified by Blobtools. using assembly_stats.py v.0.1.4 (Trizna 2020). Gene completeness was evaluated using compleasm v.0.2.5 (Huang and Li 2023) using the insecta_odb10 ortholog set.

### Whole Genome and Silk Gene Annotation

We annotated repetitive elements in each genome with Earl Grey v.4.4.0 (Baril et al. 2024). We identified putative genes with Helixer v.0.3.4 (Stiehler et al. 2021; Holst et al. 2023) via the Helixer web server (https://www.plabipd.de/helixer_main.html), Braker v.3.0.8 (Gabriel et al. 2024), and Augustus v.3.5.0 (Hoff and Stanke 2019) and selected the highest quality annotation for each genome based on the protein BUSCO completeness score. Functional gene annotation was completed by BLAST searches of protein sequences from the annotation against the non-redundant NCBI protein database and using Blast2GO v1.5.1 (Conesa and Götz 2008).

Silk *e-fibroin* sequences were annotated using a combination of *ab-initio* prediction and manual curation. Generally, *e-fibroin* paralogs were identified in full by Helixer during feature annotation. However, to corroborate true hits, putative genes were confirmed using BLASTn for terminal sequences of *e-fibroin* genes from other embiopteran species (Harper et al. 2021). For *O. nigra,* we mapped -body RNAseq reads from three individuals to each *e-fibroin* paralog and visually inspected for coverage peaks at each putative exon annotation using Geneious and IGV visualization software. For *A. ceylonica,* we used PacBio long-read IsoSeq transcriptome sequences to verify exon boundaries. For all silk genes, we confirmed the presence of signal peptides using the program SignalP v.6.0 for predicting cleavage sites. To ensure that annotated silk gene repetitive regions were not an artifact of assembly error or collapse, we mapped raw reads back to each contig containing gene clusters using minimap2, and visualized coverage peaks in IGV.

### Phylogenetic Analyses

To better understand the evolutionary history of gene duplications in the *e-fibroin* gene family, we reconstructed phylogenetic trees of the non-repetitive N- and C-terminal regions for all available embiopteran representatives (Figure 2b), as these sections of the gene are highly conserved between species. In previous studies (Collin et al. 2011; Harper et al. 2021), sequence identity and similarities were reported for partial termini sequences of a single copy of *e-fibroin* per species. However, due to differences in experimental design (e.g. sequencing cDNA vs. long-read DNA), these comparisons were conducted only using the C-termini, as this was more readily recovered across all available sequences (Fig. 3b). Here, we incorporate both N- and C-termini of all paralogs in our alignment to recover greater phylogenetic resolution, and to include multiple gene copies. To reconstruct a phylogeny using the conserved N- and C-termini dataset and the intron dataset, sequences were aligned using MAFFT v.7.490. Maximum-likelihood phylogenetic analyses were conducted in IQTree v.3.0.1, with model selection based on the Bayesian Information Criterion (BIC) in the default settings of ModelFinder. No partitioning scheme was applied, and each alignment was analyzed as a single partition. The best-fit model identified for the maximum-likelihood analysis was LG+G4. Branch support was assessed using 1,000 ultrafast bootstrap replicates. Resulting trees were visualized in FigTree v.1.4.4 and rooted using the *e-fibroin* sequence from most distantly related outgroup species within Embioptera, *Antipaluria urichi*.

## Data Availability

The Whole Genome Shotgun project for *Apothonia ceylonica* has been deposited at DDBJ/ENA/GenBank under the accession JBEUDN000000000. The version described in this paper is version JBEUDN010000000. The PacBio HiFi reads and Isoseq reads are available under BioProject PRJNA1109428. The Whole Genome Shotgun project for *Oligotoma nigra* has been deposited at DDBJ/ENA/GenBank under the accession SAMN51096571. The version described in this paper is version SAMN51096571. The PacBio HiFi reads are available under BioProject PRJNA1109428. Annotations and all other supplemental material mentioned in this manuscript can be found at the following FigShare repository: doi 10.6084/m9.figshare.30896357

## Acknowledgements

We would like to thank the staff at the DNA Sequencing Center at Brigham Young University who prepared and sequenced the embiopteran specimens used in the report, especially the late Dr. Ed Wilcox; and Dr. Riley Nelson for photographing *Aposthonia ceylonica* specimens. We would also like to thank the Institute for Comparative Genomics staff at the American Museum of Natural History for their help troubleshooting molecular lab protocols.

## Funding Sources

This work was supported by College Undergraduate Research Awards from the College of Life Sciences at Brigham Young University, the Richard Gilder Graduate School at the American Museum of Natural History, the Society for Systematic Biologists Graduate Student Research Award program, and the United States National Science Foundation’s Graduate Research Fellowship Program award 1938103 to AM and MCB awards 2217155 to PBF and 2217158 to RHB and CYH.

## Author contributions

AM, LJB, RHB, PBF conceived the study. AM, LJB, PBF secured funding. Insects were originally reared and donated by JSE who provided expertise on Embioptera. Data was analyzed and figures designed by AM, LJB, DDD, AP, RHB, PBF. CT scans were performed and formatted as figures by AM, ELS, and AYK. CYH, JLW, RHB, and PBF provided advising and funding. AM, LJB, DDD, RHB, PBF wrote and revised the manuscript. All authors edited and approved the paper.

## Conflict of Interest

The authors do not have any conflicts of interest to report.

